# Facilitated analysis of large data sets by interactive visualisation

**DOI:** 10.1101/178616

**Authors:** Zhigang Lu, Yanyan Zhang

**Author notes:** Corresponding to: Z. Lu.

## Abstract

In biological research analysis of large data sets, such as RNA-seq gene expression, often involves visualisation of thousands of data points and associated database query. Static charts produced by traditional tools lack the ability to reveal underlying information, and separated database query is laborious and involves a lot of manual effort. Interactive charting is able to make the data transparent but the use of visualisation tools often requires certain programming skills, which hinders most academic users. We present here an open-source chart editor for interactive visualisation, which is designed for academic users with no programming experience. It can not only visualise the data in an interactive way, but also link the data points to external databases, by which the user can save a lot of manual effort. We believe that interactive visualisation using such tools will facilitate analysis of large data sets as well as presenting and interpreting the data.

---

It’s the era of big data. As sequencing technology improves and the cost lowers down, more and more large data sets are generated at any time, in particular from various –omics studies (genomics, transcriptomics, proteomics, etc.). RNA-seq is one of the widely used approaches to investigate gene expression, which has evolved from sequencing of the whole organism to a single cell (scRNA-seq)^1^. As sequencing data are generated much faster than before, quick and accurate data analysis is also desirable, which involves not only statistical testing, but also gathering information from different resources to interpret the testing results. One scenario is differential gene expression analysis. Because for most eukaryotes there are at least thousands of genes^2^, one very often will have to handle a large number of data points, for which visualisation and database query are commonly needed. However, in most cases these steps are performed separately and involve a lot of manual effort. In addition, most traditional charting tools can only generate static images, from which the information on underlying data cannot be obtained, and sometimes the conclusion based on the static feature can be biased.

Interactive visualisation can facilitate the analysis of large data sets in many ways. Firstly, the data is transparent, and by interacting with the data, one can get both the overview and details about single points. Secondly, in some cases, one can also manipulate the data by customising thresholds, changing charting options and even switching data sets, which gives “multiple dimensional” sense of the data. Thirdly, by linking the data points with desired external databases or resources, the user can save a huge amount of time and manual effort. Furthermore, interactive chart is always beneficial for presenting the results, interpreting the data and discussion.

Although powerful in presenting and analysing data, interactive charting has been seldom used in scientific research, and one of the main hurdles is that most open-source charting tools require certain programming skills (e.g., Python, JavaScript or R). Nevertheless, efforts have been made to promote the transformation of images from static to interactive in scientific research^3^, including those from publishers (e.g., the Elsevier Interactive Plot Viewer^4^).

Here we would like present an interactive chart editor (IVIS) for people without any programming experience. It can be used for facilitating analysis of large data sets, for generating chart files for publication, as well as presenting the data.

IVIS uses the charting functions of HighCharts^5^ (a JavaScript library) and has the following features:

1. It’s easy to use IVIS directly handles user’s input containing comma-, tab-, or space-separated values. It can also transpose the data (i.e. switch the rows and columns) when necessary. Settings can be directly made for title, axis names, point colours, and so on (Figure 1). Commonly used chart types are covered by IVIS, including bar / column chart, dot / scatter plot, 2D scatter chart, line chart, pie chart and heat map. Exemplary interactive charts and sample data for each chart type are provided for a quick start.

**Fig.1.**
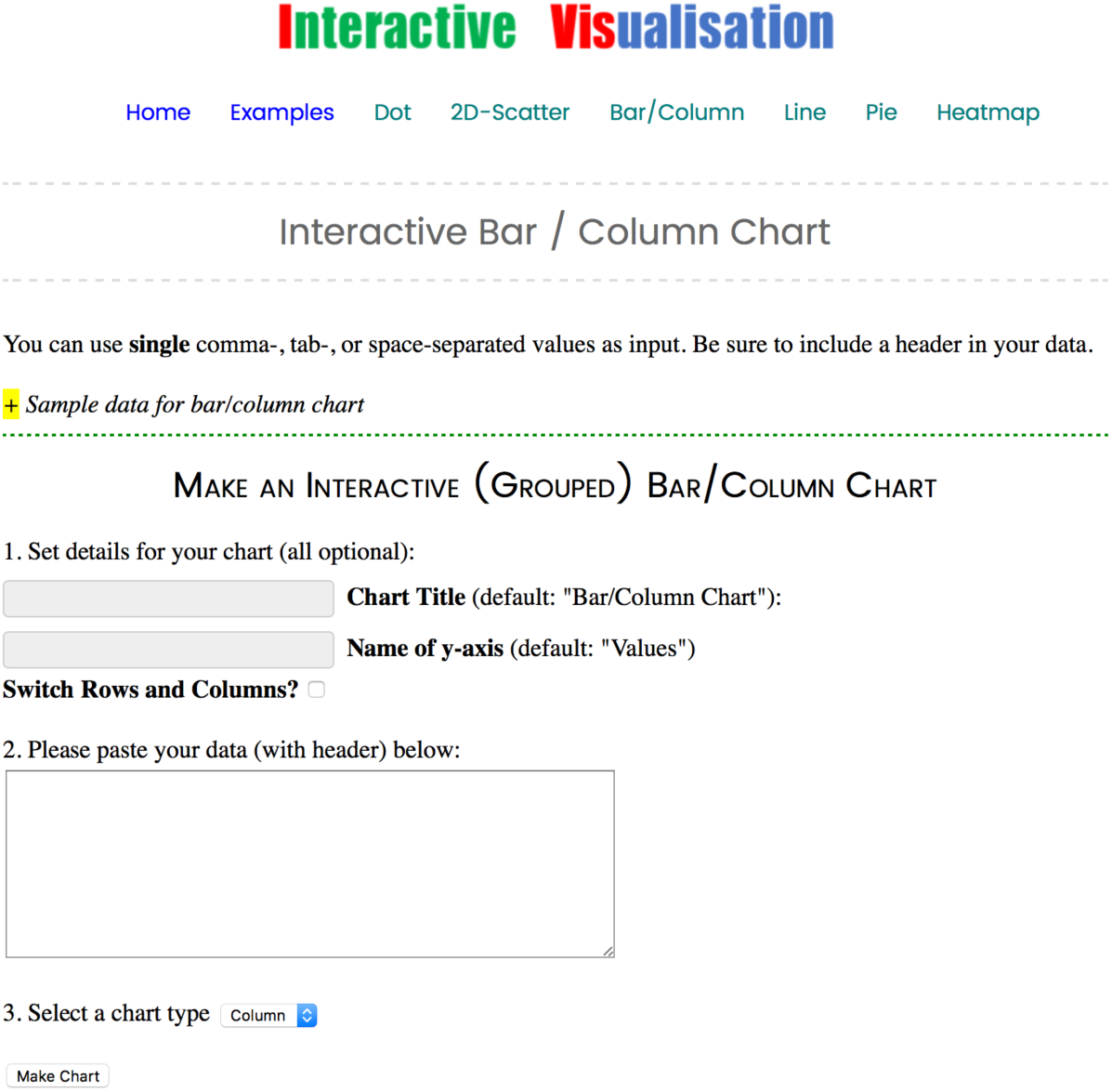
Exemplary charting interface of IVIS. Sample data are marked with ‘+’ and user can set optionally the attributes. User can switch the rows and columns when necessary.
2. The chart is fully interactive Tooltips showing data details will show up in real-time when mouse moves over. The user can zoom in at any area by simply selecting and dragging a box, and the scales will change accordingly. For multiple-series chart the user can also select to show/hide certain data. In addition, the user can customise the link for each data point, which can be an external database, the search engine, or any other file / webpage.
3. Print and export options The resulting chart can be printed, as well as downloaded it in different formats, including PNG, JPEG, PDF and SVG, at publication resolution.
4. Fast charting Thousands of data points can be plotted within a second. Tooltips and zooming are in real-time (Figure 2).

**Fig. 2.**
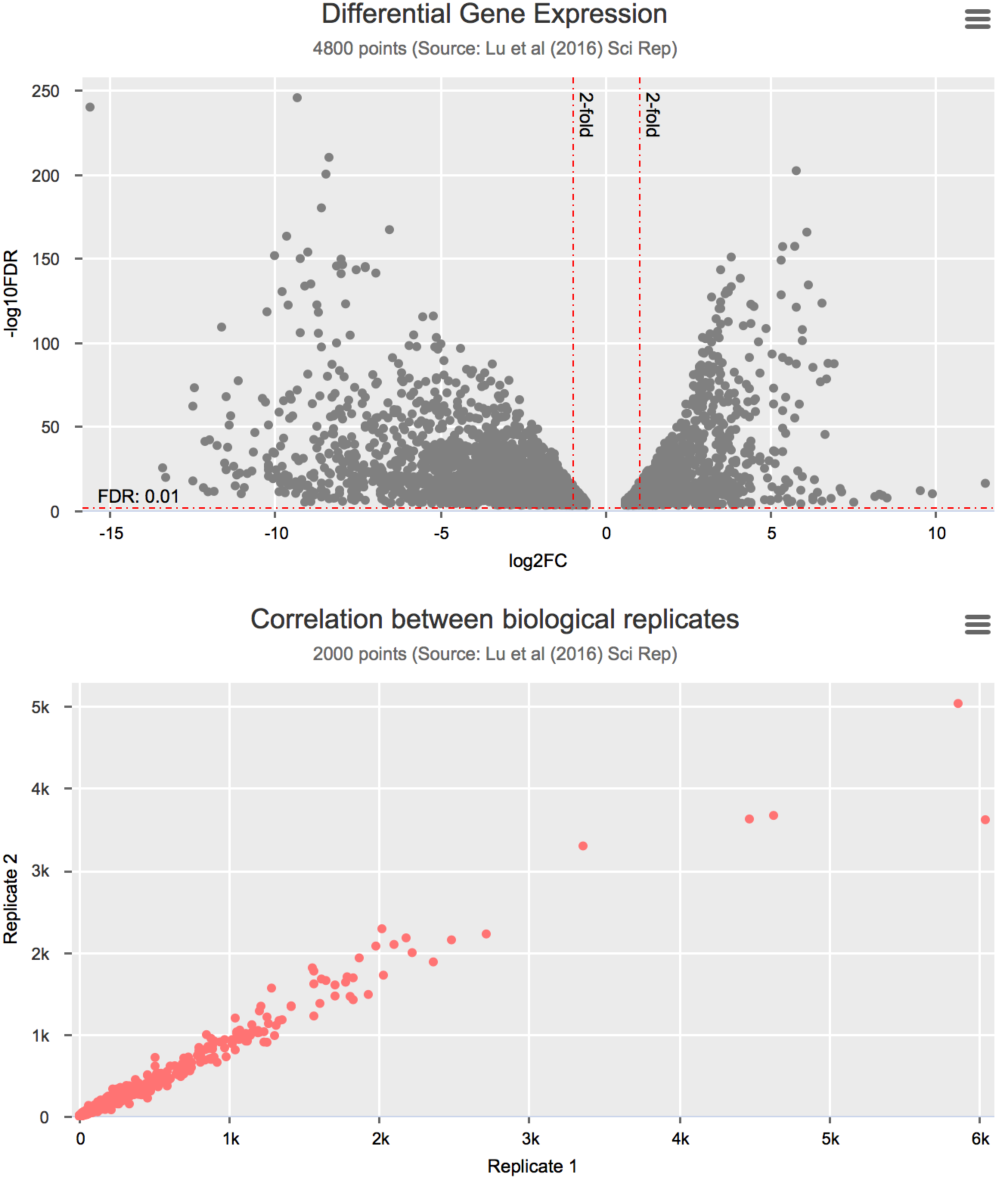
Exemplary charts. Volcano and correlation plots are widely used in RNA-seq differential gene expression analysis and very often they have thousands of data points. Here the user can link each data point with an external webpage or file to have a quick access to it
5. It can work both online and locally User can either use the online editor, or download a copy and work offline. It’s purely working on the client side (compatible with most modern web browsers).
6. It’s fully customisable The default editor includes commonly used settings. When necessary, the user can also modify the code and make their own copy.
7. It’s open-source IVIS is released under the Creative Commons license. We hope to make it better by open-sourcing the code.

## Perspective

With the presented tool we would like to draw attention from researchers who are dealing with large data sets to interactive visualisation as it can provide many speedy benefits. We also hope that developers can contribute to make interactive graphing less painful for academic users who have little programming experience.

## Availability

The code of IVIS is publicly available at the project page: <u>https://github.com/zglu/ivis.</u>The chart editor can be accessed from <u>https://ivis.xyz.</u>

